# Sub-Cluster Identification through Semi-Supervised Optimization of Rare-cell Silhouettes (SCISSORS) in Single-Cell Sequencing

**DOI:** 10.1101/2021.10.29.466448

**Authors:** Jack Leary, Yi Xu, Ashley Morrison, Chong Jin, Emily C. Shen, Ye Su, Naim Rashid, Jen Jen Yeh, Xianlu L. Peng

**Author notes:** Correspondence &.

## Abstract

Single-cell RNA-sequencing (scRNA-seq) has enabled the molecular profiling of thousands to millions of cells simultaneously in biologically heterogenous samples. Currently, common practice in scRNA-seq is to determine cell type labels through unsupervised clustering and the examination of cluster-specific genes. However, even small differences in analysis and parameter choice can greatly alter clustering solutions and thus impose great influence on which cell types are identified. Existing methods largely focus on determining the optimal number of robust clusters, which is not favorable for identifying cells of extremely low abundance due to their subtle contributions towards overall patterns of gene expression. Here we present a carefully designed framework, SCISSORS, which accurately profiles subclusters within major cluster(s) for the identification of rare cell types in scRNA-seq data. SCISSORS employs silhouette scoring for the estimation of heterogeneity of clusters and reveals rare cells in heterogenous clusters by implementing a multi-step, semi-supervised reclustering process. Additionally, SCISSORS provides a method for the identification of marker genes of rare cells, which may be used for further study. SCISSORS is wrapped around the popular Seurat R package and can be easily integrated into existing Seurat pipelines. SCISSORS, including source code and vignettes for two example datasets, is freely available at https://github.com/jrleary/SCISSORS.

## Introduction

Single-cell RNA-sequencing (scRNA-seq), which has garnered tremendous interest in the field of biomedical research in the past decade, enables the study of the transcriptome of a tissue or organ at an unprecedented resolution (1,2). Compared with bulk RNA-seq which can only capture the average of gene expression in a sample, scRNA-seq enables the profiling of the transcriptome in each individual cell simultaneously, which can theoretically lead to a more comprehensive and exhaustive understanding of each cell type. However, the computational analysis of scRNA-seq data is often challenging due to its noisy and high dimensional features (3,4).

With identifying cell populations being one of the principal goals of employing scRNA-seq, accurate analytical approaches for clustering cells are critical. To this end, a wide variety of clustering algorithms have been employed in scRNA-seq data analysis. For example, *K*-means, a popular algorithm used widely across many fields, has been implemented by methods such as GiniClust3, SC3, and RaceID (5–7). However, one of the assumptions of *K*-means clustering method is that the clusters are of similar sizes, which is often violated in the scRNA-seq data, where cell clusters are usually of widely varying sizes (8). This may negatively impact the performance of *K*-means and is exacerbated in the presence of rare cell types. Another method, Spectrum, has been created to accommodate negative-binomially-distributed single cell counts, through its implementation of spectral clustering on only 300 to 8,500 cells and 100 highly variable genes (9). However, spectral clustering is computationally expensive, with runtime scaling cubically with sample size, which is a constraint as single cell datasets have grown from thousands to millions of cells (4). Recently, the graph-based Louvain modularity optimization clustering algorithm as implemented in Seurat and Scanpy has been shown to be very accurate and achieves reasonable runtimes (10–13). The IKAP method is built around Seurat and begins by first overclustering and then transitioning to a coarser clustering by iteratively combining the clusters whose centers are closest (14). However, IKAP performance may be limited on highly heterogeneous samples such as tumors, or development datasets where expression is expected to exist on a gradient (14). Similarly, to IKAP, the recently developed MultiK method attempts to find the true number of clusters *K* in a dataset by iteratively testing many combinations of the resolution parameter *r* (15). Both IKAP and MultiK are time-intensive due to the design of iterating over parameters (14,15). In addition, SAFEclustering is an ensemble method that derives a consensus clustering from the results of several different clustering methods (16).

Besides the general demand of calling cell types in a dataset, the identification of rare cell types may be of particular interest. A plethora of clustering methods have been developed specifically for this purpose, as the identification of rare cell types from scRNA-seq data is challenging and necessitates special methodologies. For example, GiniClust3 identifies marker genes through Gini indexing, then runs DBSCAN in an attempt to find rare cell types (5). However, it uses *K*-means as part of the final consensus clustering, which may compromise the accuracy of its results as the *K*-means assumptions are often violated as previously described. This is the same case for RaceID, which was designed for rare cells but also employs the *K*-means algorithm for clustering (7).

The gap in methods for rare cell identification motivated our development of SCISSORS, which is wrapped around Seurat and strives to achieve a biologically optimal clustering. SCISSORS employs carefully designed initial clustering and reclustering steps, which ensures the identification of cells types that are of extremely low-abundance. In SCISSORS, the silhouette coefficient is used to allow the systematic estimation of intra-cluster heterogeneity, which can help determine if reclustering is needed for a cluster when biological information is insufficient. In the reclustering step, SCISSORS tests several user-defined combinations of parameters and determines the best parameter set, which helps optimize the final clusters systematically. By applying SCISSORS to a peripheral blood mononuclear cell (PBMC3K) dataset, as well as a pancreatic ductal adenocarcinoma (PDAC) dataset, we demonstrated that SCISSORS is able to identify cells that were overlooked in other analyses possibly due to their rare representation in the dataset.

## MATERIALS AND METHODS

### The SCISSORS framework

SCISSORS is a carefully designed tool wrapped around the Seurat package for the identification of rare cell types. The motivation of the methodology is rooted in the fact that cell types in a given dataset are very rarely of uniform abundance, and similarity levels among broad cell types versus among subpopulations of the same cell type can vary widely. This has made the determination of the true number of cell clusters *K* difficult, especially when cell types of low abundance exist in the dataset. To solve this problem, SCISSORS proposes to execute a multi-step process that splits potentially heterogenous cell clusters into subclusters, which leads to the identification of cell types of low abundance (Figure 1).

**Figure 1.**
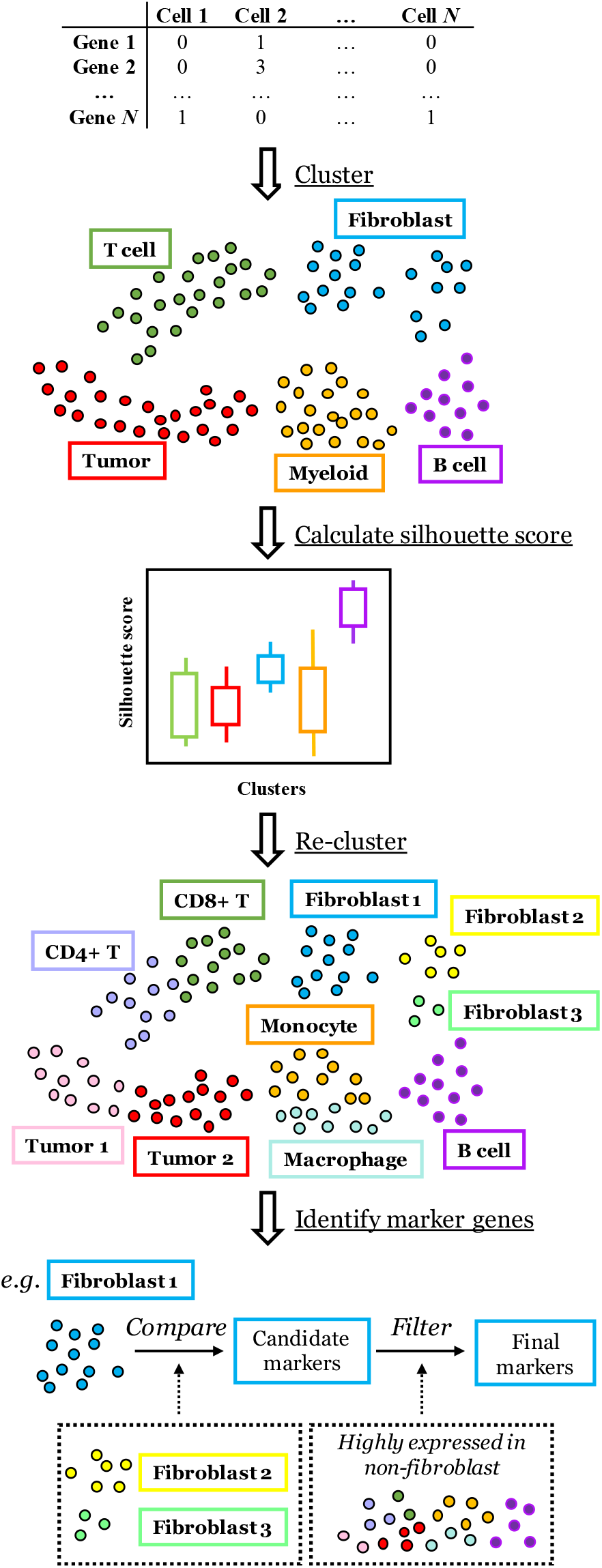
Overview of the SCISSORS framework. SCISSORS is a wrapper of the Seurat package and integrates multi-step semi-supervised clustering to perform cell type identification for even under-represented cell types in single-cell RNA-seq datasets. SCISSORS first broadly defines major clusters using Louvain clustering with conservative parameters. Then silhouette scores are calculated to infer the heterogeneity of each major cluster. Lower median silhouette score indicates higher heterogeneity of a cluster. These scores combined with biological knowledge aid the user in determining reclustering candidates. Next, combinations of reclustering parameters are tested, and the set that maximizes the silhouette scoring metric is chosen as optimal. Finally, marker genes are identified for each of the clusters and sub-clusters in order to facilitate cell type annotation and further study. For the accurate identification of marker genes in subclusters or rare cells, SCISSORS integrates a carefully designed function, where the subcluster is first compared to closely related clusters followed by a filtering step to remove highly expressed genes in the rest of the clusters.

With a pre-processed count matrix, the SCISSORS framework first performs an initial clustering step to define major clusters using conservative parameters (Figure 1). To evaluate the level of heterogeneity of each major cluster, SCISSORS calculates the silhouette score of each cell, which measures how well cells fit within their assigned clusters (17). The silhouette score estimates the relative cosine distance of each cell to cells in the same cluster versus cells in the closest neighboring cluster. Therefore, a lower median silhouette score or large variations in the distribution of silhouette scores in a cluster can be interpreted to indicate higher heterogeneity in that cluster. Then, heterogeneous major clusters suggested by silhouette scores or major clusters of biological interest can be manually selected and further considered in the reclustering step by SCISSORS. In this way, major clusters are pre-filtered for the second round of reclustering to avoid over-clustering on originally homogeneous or biologically unimportant cell clusters.

In the second round of clustering, SCISSORS re-identifies highly variable genes from all the gene features (18), considering only the selected candidate major cluster(s) instead of using the genes identified in the initial processing of the entire dataset. By doing so, SCISSORS avoids losing features that are essential in distinguishing closely related cell subpopulations from one another, but are minimally representative on the scale of the entire dataset. In addition, SCISSORS also enumerates several combinations of clustering parameters to achieve optimal performance by computing and comparing their silhouette coefficients. The identified resultant subclusters can then be integrated back into the original dataset by SCISSORS and visualized (Figure 1). Additional reclustering can also be performed on subclusters to further subdivide cell populations until the cell types are considered fully explored.

SCISSORS also integrates a function to aid with the identification of marker genes for rare cell types (Figure 1). This function first compares the rare cells to other cells that fall into the same major cluster(s) or are empirically determined as closely related cell subpopulations to derive a candidate gene list. This is because comparison within the mixture of the entire dataset may miss small differences between rare cells and their closely related cells. Then this function filters the candidate gene list by removing the highly expressed genes in all the other clusters in the dataset. By doing so, SCISSORS can accurately identify distinguishing marker genes for rare cell types.

### Initial clustering

Data preprocessing is performed using Seurat. Gene expression is normalized using regularized negative binomial regression as implemented in SCTransform (18), which also provides variance estimates for each gene. The top 4,000 highly variable genes (HVGs) are used to reduce the dimensionality of the gene-cell matrix through PCA, and the first 20 principal components are used as an initialization for a two-dimensional Fast Fourier Transform-accelerated t-SNE embedding (19,20). The cells are clustered in PCA space using Louvain modularity optimization after being embedded in a shared nearest-neighbors graph. For the PBM3K dataset, *k* = 52 (the square-root of the number of cells in the dataset) and *r* = 0.4 are used. For the PDAC dataset, *k* = 155 (the square-root of the number of cells in the dataset) and *r* = 0.4 were used.

### Silhouette score

SCISSORS uses silhouette scores (17) based on the cosine distance between cells to estimate the heterogeneity of derived clusters and to evaluate parameter sets during reclustering. To measure the dissimilarity or distance between two non-zero vectors, the cosine distance is defined as:

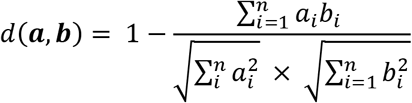

where ***a*** and ***b*** are two vectors each of length *n*. The mean distance from cell *i* to all other cells *j* in cluster *C_m_* is defined as:

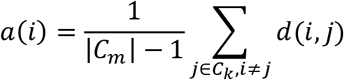

where *d*(*i, j*) is the cosine distance from cell *i* to cell *j*. The minimum mean distance from cell *i* to all points in all other clusters *C_l≠m_* is then calculated as:

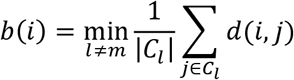

The silhouette score of each individual cell in cluster *C_m_* is then computed using the following:

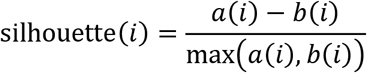

The silhouette coefficient of cluster *C_m_* is given by:

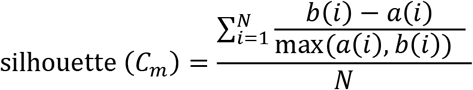

where *N* is the number of cells in cluster *C_m_*.

### Reclustering Optimization

After cells are clustered and broadly defined in the initial round of clustering, one or more rounds of reclustering are used to identify subclusters within candidate major cluster(s). The number of rounds is determined by the user based on the following two criteria. One is if the final clusters are biologically desired; the other one is if the final clusters are too homogenous to be reclustered, which can be estimated by the silhouette scores.

Rare cell types sometimes show subtle signatures from genes that are excluded in the initial round of clustering due to a low overall variance in expression. Therefore, all the genes from the major cluster(s) are re-considered, instead of only using the gene features selected in the initial clustering. Using only the selected major cluster(s), gene expression is normalized, 4000 HVGs are selected, and dimension reduction is performed similarly to the pre-processing in the initial clustering step.

Then SCISSORS iterates a user-defined set of possible values for the *k* nearest-neighbors and resolution *r* parameters, which are used to embed the cells into an SNN graph and sort them into clusters. The default setting of SCISSORS is to compare three *k* and four *r* values, leading to a total of twelve clustering results. Each result is evaluated by computing the silhouette coefficient *S* of the dataset, which is derived by averaging the silhouette coefficients of each sub-cluster:

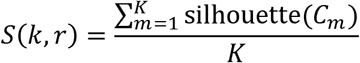

where *K* is the resultant number of clusters found using each parameter set. The set of parameters with the maximum score is then chosen as the best fit to perform the reclustering. If no reclustering results have a silhouette coefficient greater than a threshold (0.25 by default), then all reclustering results are discarded and the candidate cluster is determined to not contain further subclusters.

### Application of default Seurat, SAFE-clustering and GiniClust3 to PBMC3K

The default Seurat parameters are used to process the PBMC3K dataset as described in the Seurat v3 PBMC3K vignette (Seurat vignette) (21). Briefly, two thousand HVGs were selected using the variance-stabilizing transformation method and was used as input to PCA. The cells were then clustered in PCA space with 10 principal components using *k* = 20 nearest-neighbors and *r* = 0.5.

In SAFE-clustering, SC3, Seurat, and *t*-SNE + *k*-means are used to create an ensemble clustering, with the filtered counts matrix as input. The optional CIDR method is not used as doing so is no longer considered best practices as per the authors’ recommendation. For the SC3 clustering, the default parameters are used. The optimal value of *k* is chosen automatically, and the cells are clustered using *k*-means on the eigenvectors of the distance matrices (6). For the Seurat method, cells are clustered in 15-dimensional PCA space using *k* = 20 nearest-neighbors and *r* = 0.7. For the t-SNE + *k*-means clustering, the per-cell counts are transformed to counts per million mapped reads (CPM) and embedded in 3-dimensional *t*-SNE space with the perplexity set to 30. The optimal value of *k* is determined using adaptive density peak detection [3]. The final optimal value of *k* chosen during the ensemble clustering is determined to be equal to 9.

GiniClust3 is used to cluster the cells using the normalized and filtered counts matrix as input, as described (5,22). *k* = 7 nearest neighbors is used when clustering the cells based on their Gini indexes. Default parameters are used when clustering the cells based on their Fano factors. The consensus clustering is also generated using default parameters.

The identified clusters from different methods are annotated using the marker genes from the Seurat v3 PBMC3K vignette (21). A silhouette score for each cell in each cluster derived from each method is computed for comparison.

### Marker gene identification

Marker genes major clusters or subcluster are identified using a specifically designed integrative method, which is included as a function in SCISSORS. In this method, a given cluster or subcluster is first compared with background clusters, which may be the rest of the clusters or other closely related subclusters, to generate a candidate gene list. Subsequently, this candidate gene list is filtered by removing highly expressed genes in the clusters that are not of interest. The highly expressed genes are defined as the top expressed (top 10% by default) by averaging all the cells within each cluster; a user-defined percentile cutoff can also be supplied.

In PBMC3K dataset, for each major clusters in the first round of clustering, a candidate gene list is derived by comparing each given major cluster to the rest of the clusters (Wilcoxon rank-sum test, adjusted *p* < 0.05, log2 fold change > 0.25). This candidate gene list is then filtered by removing the highly expressed genes (top 5%) in the rest of the clusters. For the T-cell subclusters (T0.Naive CD4+ T, T1.Memory CD4+ T and T2.CD8+ T), the marker genes are derived by first comparing each of the T-cell subclusters with the rest of the T-cell subclusters (Wilcoxon rank-sum test, adjusted p < 0.05, log2 fold change > 0.5). Then the candidate gene list is filtered by removing the top 5% genes in each of the non-T-cell clusters. For the monocyte subclusters (M0.Classical monocyte, M1.Intermediate monocyte and M2. Non-classical monocyte), marker genes are derived by first comparing each of the monocyte subclusters with the rest of the monocyte subclusters (Wilcoxon rank-sum test, adjusted p < 0.05, log2 fold change > 0.5). Then the candidate gene list is filtered by removing the top 5% genes in each of the non-monocyte clusters.

In the PDAC dataset, for each major clusters in the first round of clustering, a candidate gene list is derived by comparing each given major cluster to the rest of the clusters (Wilcoxon rank-sum test, adjusted *p* < 0.05, log2 fold change > 0.1). This candidate gene list is then filtered by removing the highly expressed genes (top 10%) in the rest of the clusters. For the basal-like cluster, basal-like cells are first compared with the rest of tumor cells (classical 1 and classical 2) (Wilcoxon rank-sum test, adjusted p < 0.05, log2 fold change > 0.25). Then the candidate gene list is filtered by removing the top 10% genes in each of the non-tumor clusters. For the classical 1 and classical 2 genes, similar method is used. For apCAF genes, apCAF cells are first compared with the rest of the fibroblast cells (iCAF and myCAF) (Wilcoxon rank-sum test, adjusted p < 0.05, log2 fold change > 0.25). Then the candidate gene list is filtered by removing the top 10% of highly-expressed genes in each of the non-fibroblast clusters. For the iCAF and myCAF genes the same method is used.

### Bulk RNA-seq analysis

To call tumor subtypes in the bulk dataset, PDAC classical and basal-like genes are identified using the SCISSORS function. Specifically, while the basal-like gene list is derived the same way as mentioned before, the classical 1 and classical 2 cluster are combined to compare with the basal-like cluster. Subsequently, top 10 genes for the basal-like and classical cells respectively ranked by their fold-change are retained for further analysis.

The TCGA dataset is processed for the Moffitt schema and PurIST calls as described before (23,24). Unsupervised consensus clustering (kmeans) is applied on the distance matrix (Pearson) of the log2 transformed data using top 10 SCISSORS derived tumor genes (ConsensusClusterPlus version 1.56.0). Survival of subtypes is analyzed by log-rank test.

### Data access

The PDAC scRNA-seq data from the study by Elyada et al. can be obtained on NCBI dbGaP (accession number phs001840.v1.p1). The 10X Genomics PBMC3K dataset can be obtained using the SeuratData R package. TCGA normalized RNA-seq gene expression data can be obtained from the Broad Institute FIREHOSE portal [http://gdac.broadinstitute.org].

## Results

### Application of SCISSORS to the PBMC3K dataset

We applied SCISSORS to the widely used 10X Genomics PBMC3K dataset, which consists of 2,700 peripheral blood mononuclear cells from a single healthy donor (25). This dataset was previously analyzed as an example of the usage of Seurat (v3) in a vignette published by the Satija Lab (Seurat vignette) (21) in which cells were processed, clustered, and annotated. We used SCISSORS to perform an initial clustering, which yielded 6 clusters (Figure 2A) with varying silhouette scores (Figure 2B) and marker genes (Supplementary Figure S1). These major clusters were subsequently named with a prefix “P”, followed by a random cluster number. Clusters P3 and P5 were annotated as being composed of B cells and natural killer (NK) cells respectively (Figure 2A, Supplementary Figure S1), and exhibited high silhouette scores, suggesting that they were relatively homogeneous (Figure 2B). Clusters P0 and P2 were annotated as CD4+ and CD8+ T cells separately, which were more heterogeneous as defined by lower silhouette scores (Figure 2B). This, along with their biological similarity, led us to combine clusters P0 and P2 for reclustering using SCISSORS. SCISSORS reclustering found three total subclusters (Figure 2C), which were named with a prefix “T”, followed by a random subcluster number. By analyzing their marker genes, clusters T0, T1 and T2 were annotated as Naive CD4+ T cells, Memory CD4+ T cells, and CD8+ T cells respectively (Figure 2D). These three cell types accurately reproduced the annotations in the aforementioned Seurat vignette (21). This demonstrated that the multi-step clustering method of SCISSORS was able to accurately call cell types, circumventing the need to find single values *k* and *r* that are optimal for the entire dataset, a challenging step in scRNA-seq analyses.

**Figure 2.**
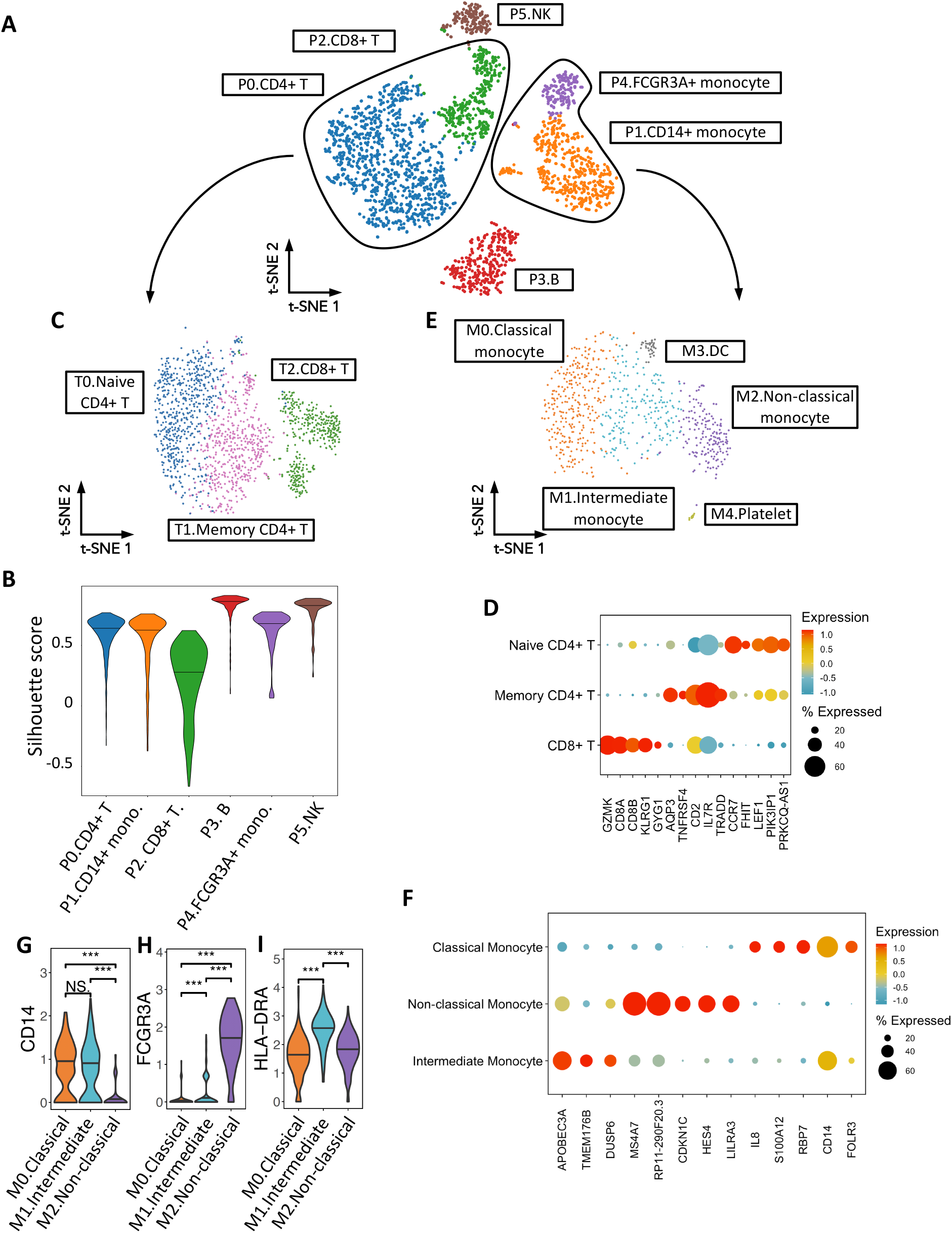
Application of SCISSORS to the PBMC3K dataset. **(A)** Initial conservative clustering on the PBMC3K dataset. Six major clusters were identified. **(B)** Violin plot for silhouette scores in each major clusters. **(C)** Reclustering of major clusters containing CD4+ and CD8+ T cells. **(D)** Top 5 marker genes identified by SCISSORS, ranked by fold-change. **(E)** Reclustering of candidate major clusters containing CD14+ and FCGR3A+ monocytes. **(F)** Top 5 marker genes identified by SCISSORS, ranked by fold-change. **(G), (H) & (I)** Marker gene expressions of monocyte subtypes in the three monocyte subclusters. ***: p< 0.05, Wilcoxon rank-sum test.

Clusters P1 and P4 were both annotated as containing myeloid cells and were subjected to a reclustering step as well (Figure 2A, Supplementary Figure S1). As a result, 5 subclusters were identified, named with a prefix “M”, followed by a random subcluster number (Figure 2E). Among them, SCISSORS reproducibly defined subclusters M3 and M4 as dendritic cells (DC) and platelets as in Seurat vignette (Figures 2F). The remaining cells were monocytes, which were reclustered into three subclusters (M0, M1 and M2) by SCISSORS. Human monocytes are known to be divided into three major populations, namely classical, non-classical and intermediate, which are primarily distinguished by different levels of CD14 and CD16 (FCGR3A) (26,27). Among the three monocyte subclusters identified by SCISSORS, subcluster M2 was associated with the FCGR3A+ monocytes in the Seurat vignette. It showed highest level of FCGR3A (CD16) and decreased level of CD14, which was aligned with known non-classical monocyte features (Figures 2G&H) (26,27). Therefore, subcluster M2 was annotated as non-classical monocytes in our study. Interestingly, among the cells that were annotated as CD14+ monocytes in the Seurat vignette, SCISSORS identified a subcluster M1, which showed a similarly high level of CD14 (Figure 2G) with subcluster M0, but an intermediate level of CD16 (FCGR3A) compared with subclusters M0 and M2 (Figure 2H), indicating that this may be an intermediate monocyte cluster. Additionally, M1 showed a significantly higher level of HLA-DR than the rest of the monocyte subclusters (Figure 2I), confirming that it consisted of intermediate monocytes. Therefore, we annotated subclusters M0, M1 and M2 as classical monocyte, intermediate monocyte and non-classical monocyte accordingly (Figure 2E). To our knowledge, the intermediate monocyte cluster was not identified by either the Seurat vignette or other analyses of this dataset. This demonstrated that SCISSORS is able to identify cells that may be overlooked by traditional clustering methods due to their low abundance or high degree of similarity to neighboring cell clusters. The annotated subclusters were integrated with the un-reclustered major clusters, all of which were visualized on the original t-SNE embedding (Figure 3A).

**Figure 3.**
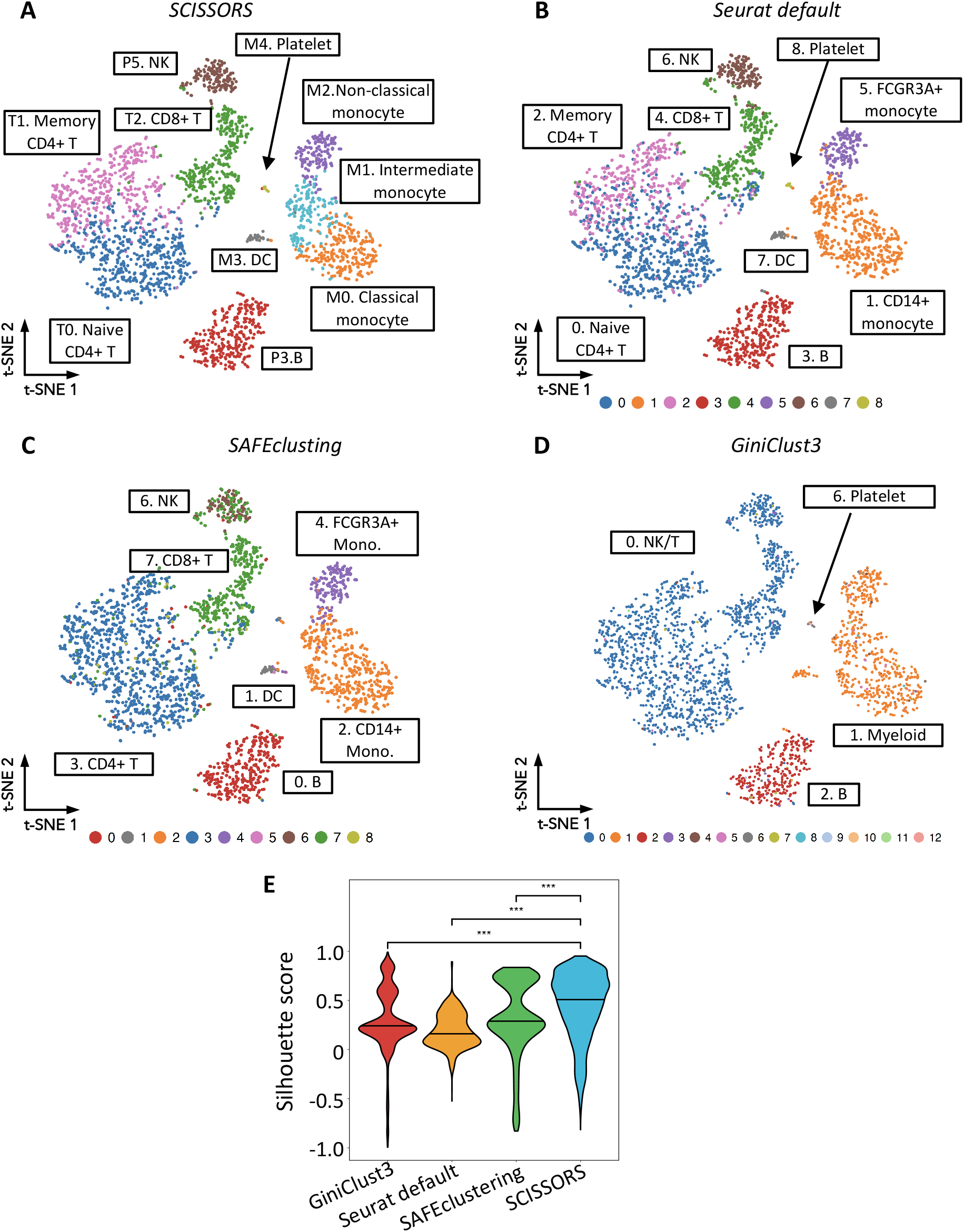
Comparison of SCISSORS to alternative methods. **(A)** Final clusters, with major clusters without reclustering and subclusters derived by reclustering in SCISSORS. **(B)** Clusters labeled by annotations derived by Satija lab analysis using default Seurat. t-SNE coordinates were derived in SCISSORS analysis in (A). **(C) & (D)** Clusters identified by using existing clustering methods SAFEclustering and GiniClust3 respectively. t-SNE coordinates were derived in SCISSORS analysis in (A). **(E)** Violin plots of silhouette score distributions of cells in the assigned clusters by each method tested. ***: p< 0.05, Wilcoxon rank-sum test.

### Comparison of SCISSORS to alternative methods

To evaluate SCISSORS against other comparable methods, we re-analyzed the PBMC3K dataset using Seurat with default parameters (Seurat default), SAFEclustering and GiniClust3 (see Methods) (5,11,16,21). Cell clusters in each method were annotated by examining the marker genes and visualized on the same t-SNE embedding obtained from our previous analysis using SCISSORS (Figures 3B-D). As already shown, compared with default Seurat, SCISSORS reproducibly found the same 8 final clusters, while also identified an extra cluster of intermediate monocytes, M1.Intermediate monocyte (Figures 3A&B). With suggested parameters, SAFEclustering didn’t separate the CD4+ T cells into their naïve and memory subtypes (Figure 3C), and also failed to cluster platelets and DCs accordingly as they were of relatively lower frequency in this dataset. In addition, SAFEclustering clustered 33 cells into a cluster (cluster 8) that we could not annotate using the Seurat vignette labels, which were scattered throughout other clusters on the t-SNE embedding (Figure 3C). Clusters identified by GiniClust3 were even more disparate, with only 4 out of 13 clusters matching Seurat vignette labels (Figure 3D). The remaining 9 clusters with 117 total cells were scattered throughout the t-SNE embedding and did not correspond to any cell types annotated in the Seurat vignette or in our analysis (Figure 3D).

Finally, to evaluate if each cluster has been sufficiently divided to reveal heterogeneous cell populations, silhouette scores for each cell corresponding to their assigned clusters were calculated for the results from Seurat default, SAFEclustering, GiniClust3 and SCISSORS. SCISSORS showed significantly higher median silhouette scores compared with each one of the other methods (Figure 3E). This demonstrated that the SCISSORS-identified clusters were of lower heterogeneity and more meaningful due to its extensive grouping of cells within major clusters into smaller, biologically relevant subclusters. In addition, SCISSORS also out-performed SAFEclustering and GiniClust3 by using significantly less runtime.

### Pinpointing overlooked basal-like cells in a PDAC dataset

Evidence has accumulated revealing the co-existence of sub-populations of tumor and fibroblast cells in PDAC patient samples (28–30). We hypothesized that SCISSORS would be able to identify these rare sub-populations even when they are extremely under-represented. Therefore, we applied SCISSORS to a published PDAC dataset from a study by Eylada et al. with 6 PDAC tumor samples and 2 adjacent normal samples (31). The initial clustering yielded 13 clusters with varying silhouette scores (Supplementary Figures S2A, B), indicating varying levels of intra-cluster heterogeneity. To gain better biological understanding and thus determine which major clusters were suitable candidates for subsequent reclustering, we used SingleR (32) to generate general cluster annotations based on references of a human primary cell dataset (33) and a human normal pancreas dataset (34). SingleR annotation revealed expected cell clusters, including epithelial/ductal cells, activated stellate/mesenchymal stem cells, acinar cells and multiple types of immune cells (Supplementary Figures S1C&D). In addition, marker genes were identified for each major cluster and overlapped with cell markers from the original study (Supplementary Figure S2E), which confirmed the general annotations from SingleR. Although this initial round of annotation was not of high enough resolution to exhaustedly determine all possible cell populations, the silhouette score distributions and annotations for each cluster allowed us to systematically determine candidates for reclustering analysis using SCISSORS.

PDAC tumors can show the presence of both basal-like and classical tumor cells, which correspond to the two intrinsic tumor subtypes found in PDAC patients (28,35–37). To understand the heterogeneity within tumor cells and study the sub-populations, we combined clusters of epithelial cells (cluster 5 and 9) (Supplementary Figure S2A) and reclustered them. This resulted in the identification of 7 subclusters (Figure 4A). The marker genes for each cluster were identified, which were compared with cell type markers identified in the original study (31) (Figure 4B). We were able to annotate five of them as Acinar, Classical 1 (two subclusters were merged in this cluster), Classical 2, Lipid processing and Secretory. In the previous study, Elyada et al. identified those same clusters, but did not detect any basal-like cells (Supplementary Figure 3A) (31). To annotate the additional sub-cluster in SCISSORS analysis, we performed scRNA-seq-based copy number variation (CNV) analysis (38). We found that this additional cluster showed strong evidence for aberrant CNVs (Figure 4C), revealing that they may be malignant tumor cells. Interestingly, by using VAM (39), we found that cells in this cluster were significantly enriched for Moffitt basal-like genes, but were absent of Moffitt classical genes (Figures 4D&E). This indicated that the cluster may consist of PDAC basal-like cells. We then used DECODER to calculate the basal-like and classical weights and the ratio between them (bcRatio) for each cell (23). DECODER weights are continuous and measure the extent of basal-like-ness and classical-ness. Interestingly, this newly identified cluster showed high basal-like weights and bcRatio, but low classical weights, confirming that this cluster may be a basal-like cell cluster (Figures 4F-H). Additionally, GATA6, which is thought to be a discriminating classical subtype marker, was found to be depleted in this cluster (Figure 4I) (40). This newly identified basal-like cell cluster was composed of only 148 cells, comprising just 0.59% of the entire dataset. Therefore, this demonstrated that SCISSORS successfully identified the basal-like tumor cells, which were rare in the dataset and were not detected in the original study.

**Figure 4.**
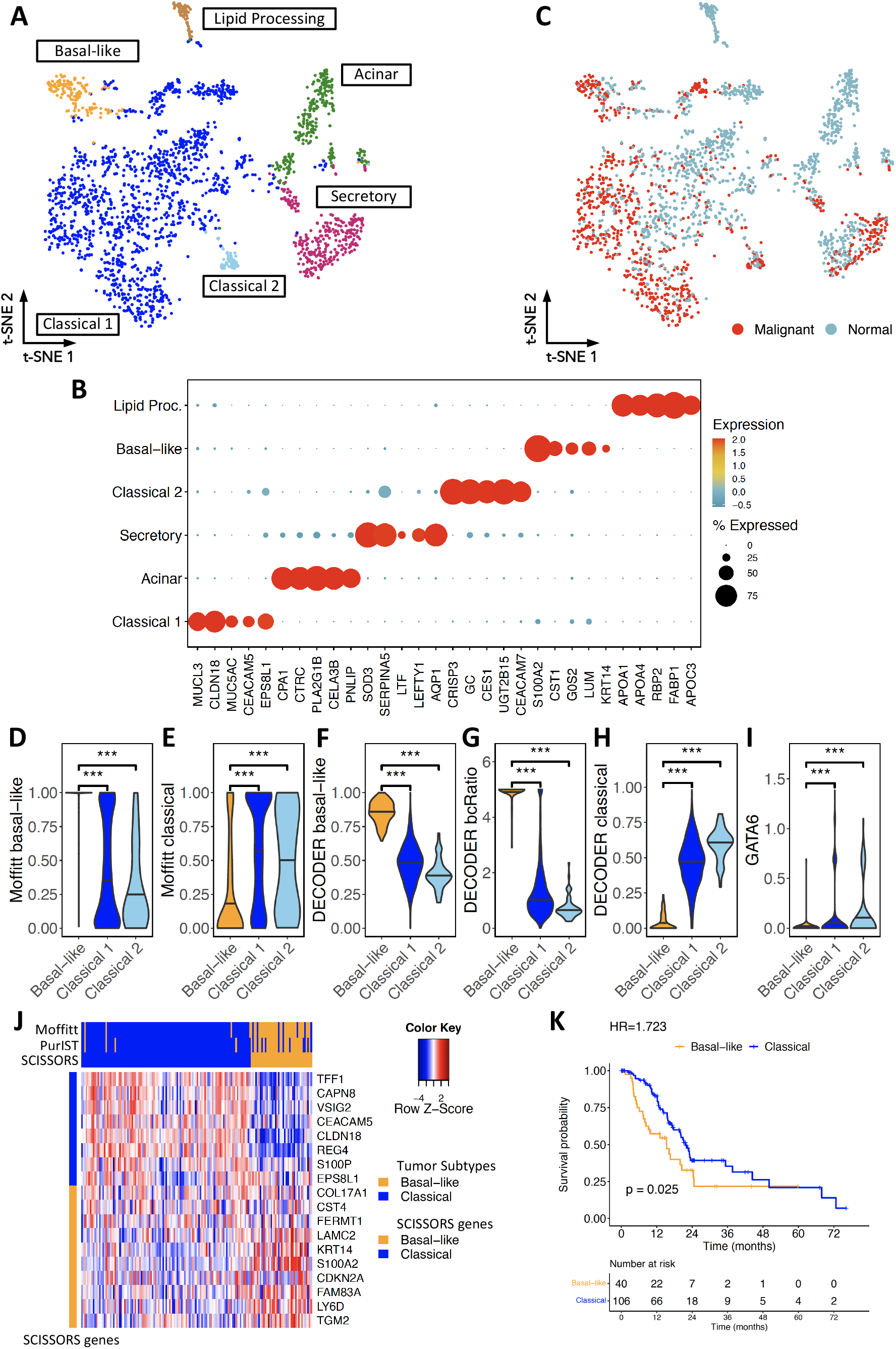
Identification of overlooked basal-like cells in the Elyada pancreatic ductal adenocarcinoma (PDAC) dataset. **(A)** Reclustering of the epithelial/ductal cell clusters revealed 6 subclusters. **(B)** Copy number variation (CNV) analysis for the identification of malignant cells. **(C) & (D)** Gene enrichment analysis for each cell in the three tumor subclusters, using VAM scores based on top 25 Moffitt basal-like and top 25 Moffitt classical genes. The identified basal-like cluster showed significantly higher basal-like gene enrichment, but lower classical gene enrichment. **(E), (F) & (G)** Tumor subtype analysis in the three tumor subclusters, using DECODER basal-like weights, the ratio between basal-like and classical weights (bcRatio) and classical weights. The identified basal-like cluster showed enriched basal-like weights and bcRatio. **(H)** GATA6 expression in the identified tumor subclusters. The identified basal-like cluster showed depleted GATA6 expression. **(I)** Top 5 marker genes identified by SCISSORS, ranked by fold-change. **(J)** Associations of tumor subtypes called by using SCISSORS derived tumor genes, with calls derived by using the Moffitt schema and the PurIST classifier. Heatmap shows consensus clustering of The Cancer Genome Atlas (TCGA) pancreatic adenocarcinoma (PAAD) samples using top 10 SCISSORS tumor genes, ranked by fold-change. **(K)** Kaplan–Meier plots of overall survival in patients with resected PDAC for tumor subtypes derived by using SCISSORS marker genes. Log-rank test was used for survival analysis. ***: p< 0.05, Wilcoxon rank-sum test.

To investigate the potential usage of scRNA-seq based and SCISSORS derived genes for analyzing bulk RNA-seq samples, we identified the top 10 basal-like and classical genes. By using The Cancer Genome Atlas (TCGA) pancreatic adenocarcinoma (PAAD), we found that the patients were clustered into the two reproducible basal-like and classical subtypes (Figure 4J). Compared with previous calls by the Moffitt schema and the PurIST classifier (24), SCISSORS based calls only showed 11 (4 basal-like and 7 classical) and 15 (4 basal-like and 11 classical) inconsistent calls out of 150 samples respectively. Additionally, basal-like patients subtyped by the SCISSORS based genes showed significant worse survival (p=0.025, Hazard ratio=1.723, log-rank test), aligned with previous findings (Figure 4K). Collectively, these demonstrated that marker genes derived from SCISSORS can be readily used to deconvolve bulk RNA-seq samples, based on the fact that cell types from SCISSORS can be highly purified and marker genes from SCISSORS are carefully filtered for specificity in the respective cell type.

### Pinpointing overlooked apCAF cells in the PDAC dataset

In PDAC, cancer associated fibroblasts (CAFs) have been shown to be heterogenous (31,41,42). In the same PDAC dataset, we applied SCISSORS on cluster 8, denoted as activated stellate/mesenchymal stem cells by SingleR, to identify CAF subtypes within the larger fibroblast population (Supplementary Figure S2A). Unlike the original study, the endothelial and perivascular cells were clustered within our initial activated stellate/mesenchymal stem cell group (Figure 5A, supplementary Figure S2A) likely due to the conservative parameters we used in the initial clustering step. In addition to the endothelial and perivascular cells, we derived three additional subclusters, all of which showed enriched pan-CAF gene expression (Figure 5B). Among them, two of the subclusters exhibited strong enrichment of inflammatory CAF (iCAF) and myofibroblastic CAF (myCAF) genes respectively (Figures 5C&D), which aligned with the discovery of iCAF and myCAF populations in the original study (39). Interestingly, marker genes for the novel cluster identified by SCISSORS exhibited a large overlap with the antigen-presenting CAF (apCAF) subtype described by Elyada et al. (31) (Figures 5E). After running an enrichment analysis of the originally defined apCAF marker genes using VAM (39), we saw that cells in this cluster showed significant enrichment (Figures 5F), confirming their apCAF identity. In the original study by Elyada et al., the apCAF population was only explicitly defined in scRNA-seq samples from KPC mouse models. Although the existence of the apCAF subtype was validated through mass cytometry staining of human PDAC sections, they were not identified in the original human PDAC scRNA-seq data analysis (Supplementary Figure 3B). Re-analysis using SCISSORS of the same human PDAC scRNA-seq data was able to identify this rare apCAF population; the apCAF cluster contained 23 cells and made up only 0.092% of the entire dataset. Thus, we again demonstrated that SCISSORS enabled the identification of extremely low abundance cells such as apCAFs, which were not identified in the original study.

**Figure 5.**
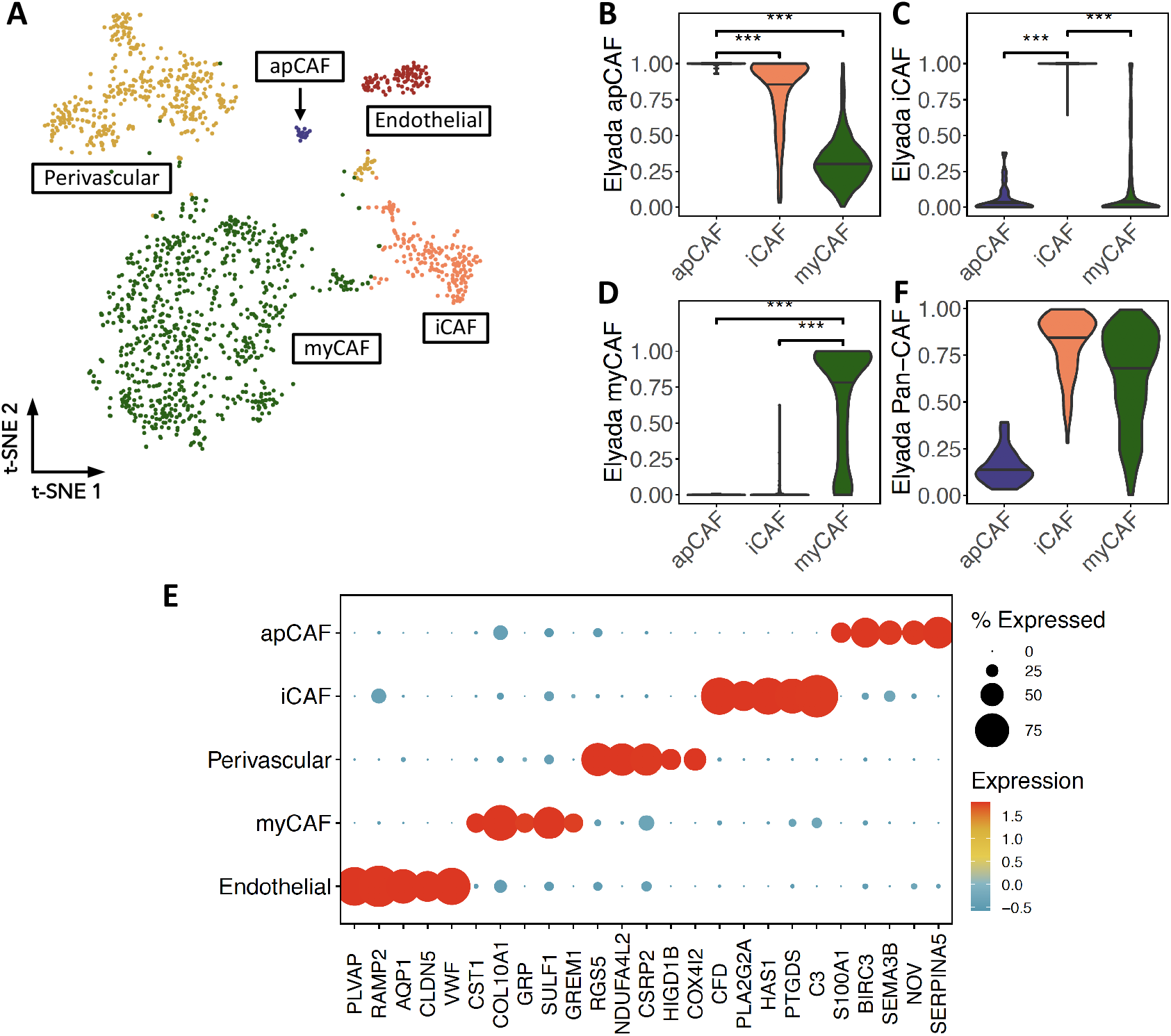
Identification of overlooked apCAF cells in the Elyada PDAC dataset. **(A)** Reclustering of the fibroblast cell clusters revealed 5 subclusters. **(B), (C), (D) & (F)** Gene enrichment analysis for each cell in the three fibroblast subclusters, using VAM scores based on Elyada pan-CAF, iCAF, myCAF and apCAF genes. An apCAF cell subcluster was identified showing apCAF gene enrichment. **(E)** Top 5 marker genes identified by SCISSORS, ranked by fold-change. ***: p< 0.05, Wilcoxon rank-sum test.

## Discussion

The optimization of clustering methods is being intensively studied in the field of scRNA-seq analysis, as it is one of the key determinants of the accurate identification of cell types through cluster-based analysis. The identification of rare cell types is even more challenging, and there are few available tools tailored to this purpose that are both accurate and efficient. To fill this gap, SCISSORS was developed based on the rationale that reclustering of heterogenous clusters can reveal rare cells.

Graph-based clustering methods such as Louvain and Leiden clustering have enjoyed growing popularity recently due to their speed and ease of use and are integrated into popular scRNA-seq analysis frameworks such as Seurat and Scanpy (11–13,43). Most clustering methods, whether graph-based or not, perform a single clustering step on the entire dataset at once, seeking to estimate parameters that are optimal for the detection of all cell types and subtypes present in the data simultaneously. In the case of Louvain modularity optimization, which SCISSORS uses, this means determining a single set of values of the number of nearest neighbors *k* and resolution *r* that both keep large and homogeneous clusters intact and separate out rare cells that might have only faint differences in gene expression from related cell types. Decreasing *k* or increasing *r* generally results in the identification of rare cells at the expense of the overclustering of truly homogeneous groups into spurious or biologically uninteresting clusters. The inverse also occurs; increasing *k* or decreasing *r* usually leads to the preservation of homogeneous clusters along with the grouping of rare cell types into closely neighboring clusters. The application of SCISSORS eliminates the need to determine universally optimal values of *k* and *r* by providing an objective and systematic measure through the silhouette score and allowing the user the flexibility to determine if each cluster is heterogenous enough to necessitate reclustering.

SCISSORS also attempts to generate a more biologically relevant clustering by re-selecting HVGs within each cluster. Rare cells appear infrequently among an ocean of other cell types, and their marker genes may be present as faint signals that might be excluded at the very beginning of an analysis due to a lack of expression in the majority of the cells and thus a low overall variance in expression. For example, in the study by Elyada et al., only 1,000 HVGs from the whole dataset were included even for subsequent reclustering analysis (31). This may be the reason why the basal-like tumor cells and apCAF stroma cells were not identified in the original analysis. To address this issue, SCISSORS reconsiders all detected genes when performing reclustering in order to include genes that may be high variance within a major cluster but are not considered as HVGs in the initial round of clustering.

To our knowledge, R is currently the most widely used software for the analysis of scRNA-seq data. Like other packages that build on Seurat, SCISSORS is freely available as an open-source R package and can be downloaded from GitHub.

## Data Availability

SCISSORS, including source code and vignettes for two example datasets, is freely available at https://github.com/jrleary/SCISSORS.

## Acknowledgements

We would like to thank the University of North Carolina at Chapel Hill and the Research Computing group for providing computational resources and support that have contributed to these research results. We would like to thank Dr. Hildreth Robert Frost (Department of Biomedical Data Science, Geisel School of Medicine, Dartmouth College) for his tremendous help in VAM package usage and result interpretation. We would like to thank Dr. Yuchen Yang (Department of Pathology and Lab Medicine, University of North Carolina at Chapel Hill) for his generous comments in best practices when using SAFEclustering.

## Author contributions

X.L.P. and J.J.Y. designed the study. J.L., Y.X, A.M., J.C., Y.S., E.C.S. and X.L.P. collected the data and performed the analyses. X.L.P., J.L., and J.J.Y. wrote the paper. All authors critically reviewed and commented on the paper.

---

**Supplementary Figure 1.**
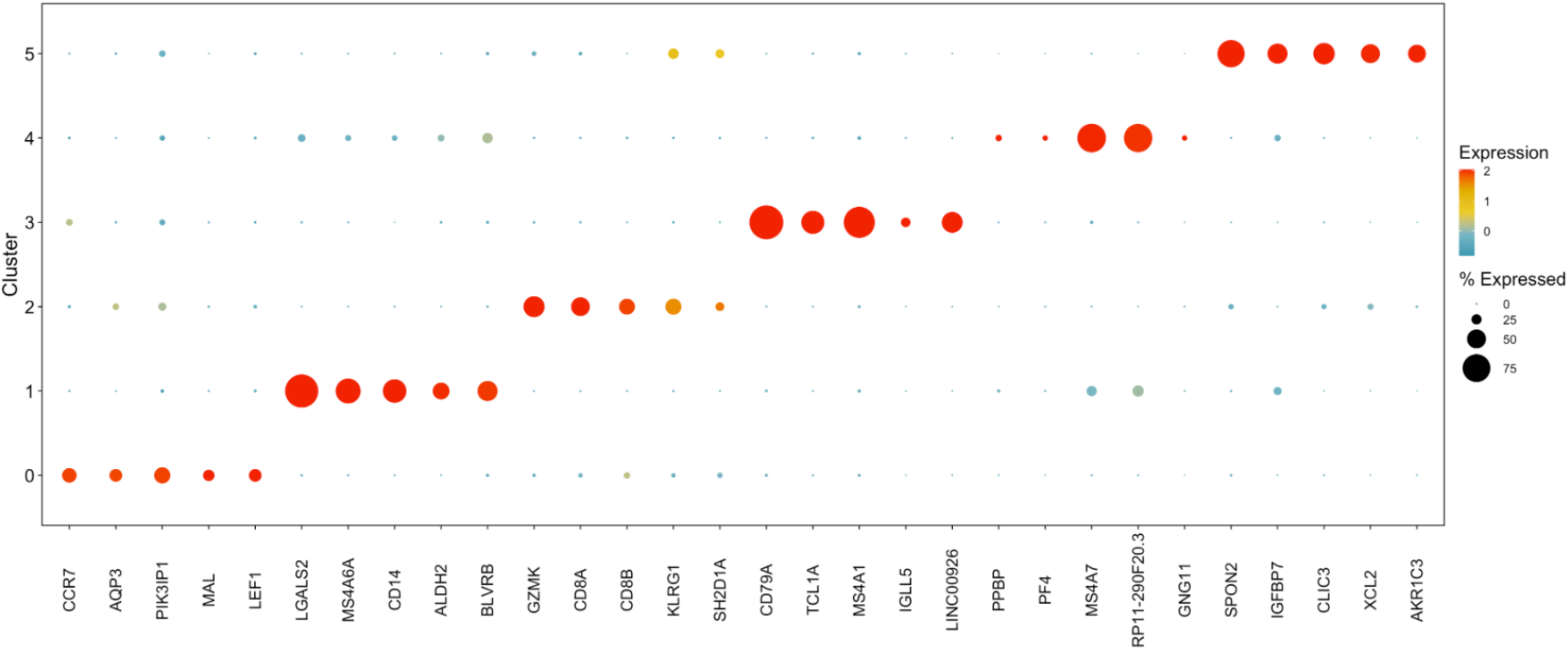
Top 5 marker genes ranked by fold-change for the major clusters identified by SCISSORS in the PBMC3K dataset.

**Supplementary Figure 2.**
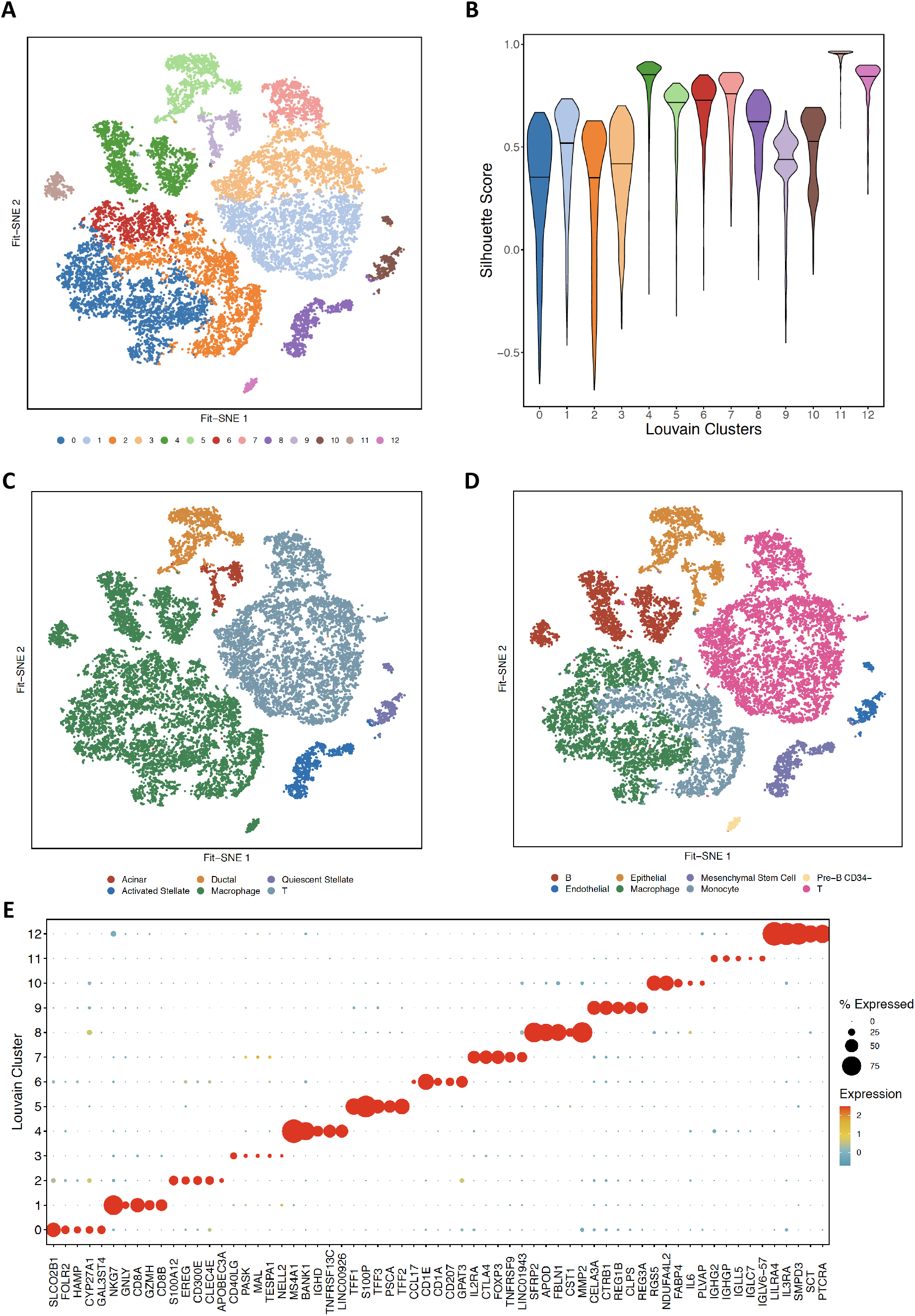
Application of SCISSORS to the Elyada PDAC dataset. **(A)** Initial conservative clustering. Twelve major clusters were identified. **(B)** Violin plot for silhouette scores in each major clusters. **(C) & (D)** SingleR annotations on the major clusters using the single-cell PDAC dataset and bulk human cells as references. **(E)** Top 5 marker genes for the major clusters identified by SCISSORS, ranked by fold-change.

**Supplementary Figure 3.**
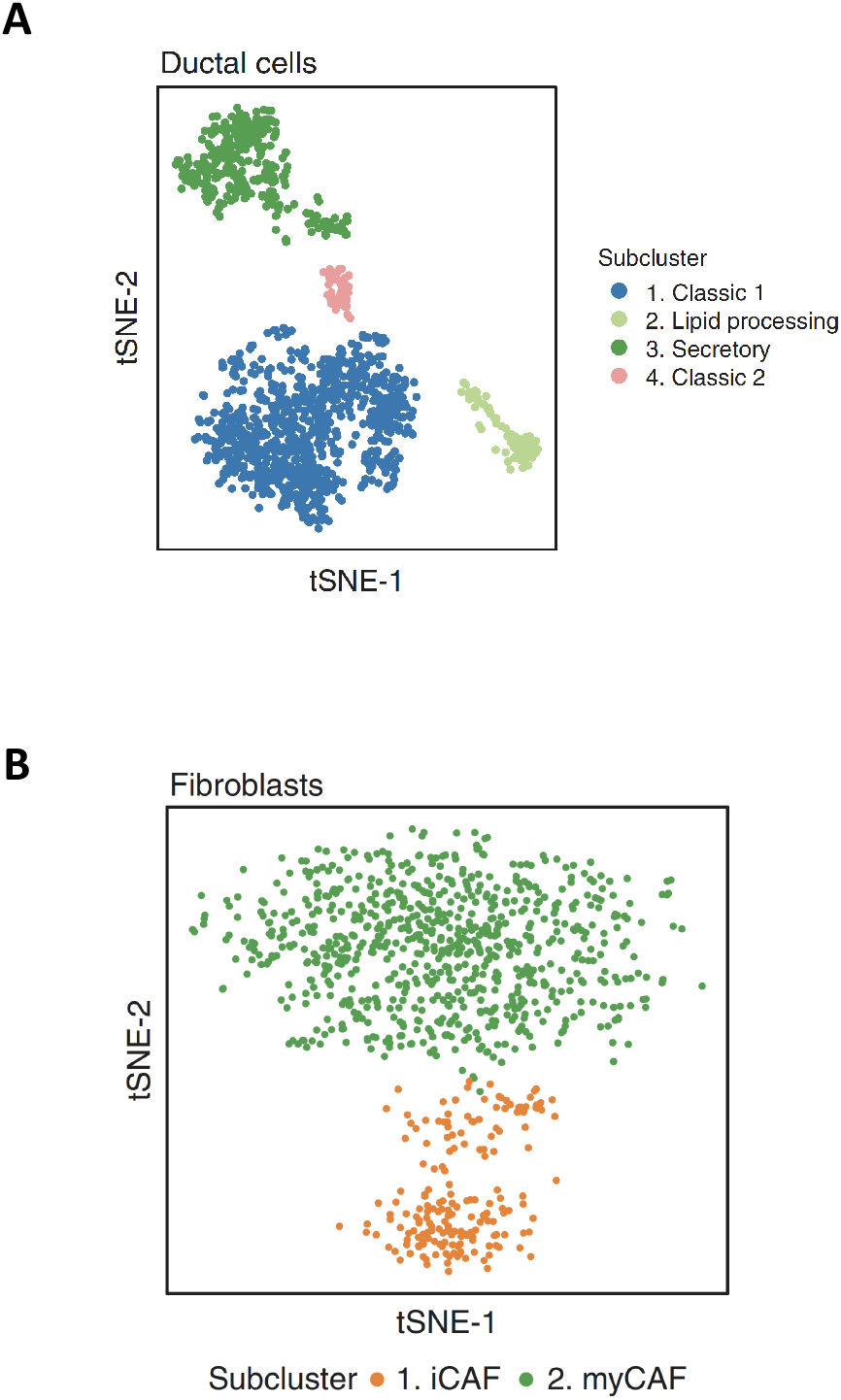
Tumor and fibroblast clusters identified in the original Elyada study. **(A)** No basal-like cluster was identified. **(B)** No apCAF cluster was identified. (Figures adapted from Figure 1D and Figure 3A in Elyada et al., *Cancer Discovery*. 2019 Aug;9(8):1102-1123., for easy comparison for the reviewers only. Copyright belongs to the authors and *Cancer Discovery*.)

* For reviewers only.

* Copyright belongs to Elyada et al. and *Cancer Discovery*.

